# Comparative genome analyses suggest a common blueprint for obligate endoparasitism in Strepsiptera

**DOI:** 10.64898/2026.09.16.752099

**Authors:** Rebecca Jean A. Millena, Paul B. Frandsen, Marisano James, Ethan R. Tolman, Floria M. K. Uy, Jessica L. Ware

## Abstract

The twisted-wing parasites (order Strepsiptera) are well-known for their unusual life histories as obligate endoparasites of a wide variety of other insect orders. Here, we present the first high-quality genome assembly of a leafhopper-parasitizing strepsipteran species. We sequenced the genome of *Elenchus koebelei* using PacBio HiFi technology, yielding a primary assembly of 63.7 Mb in 96 contigs, with an N50 of 1.60 Mb and a BUSCO completeness score of 82.3%. In total, we recovered 8,167 protein-coding gene annotations, and 34.2% of the genome annotated as repeats. For the first time, we compare the genome content of three strepsipteran species and find that they are remarkably consistent in broad structure, with approximately 600 insect universal single-copy orthologs that appear to be completely missing from the order. These similarities imply a conserved genomic blueprint for strepsipteran parasitic strategy and/or differences that must be investigated at the post-translational level.

## Background and Summary

The order Strepsiptera comprises a small group of obligate insect endoparasites, with approximately 630 described species distributed across 14 extant families and a remarkably diverse host range (Kathirithamby 2025; Millena et al. 2025). Strepsiptera are entirely parasitic, with a life cycle defined by extreme sexual dimorphism. Free-living, winged males survive only hours as reproductive adults, while females of most families are neotenic (larva-like) and remain permanently embedded within their host (Kathirithamby 2025). Until recently, baseline biological questions about Strepsiptera proved difficult to resolve (Millena et al. 2025), but genomic, transcriptomic, and phylogenetic studies have steadily expanded molecular sampling across the order (Balzer et al. 2020; Dong et al. 2022; Hui et al. 2023; Castaño et al. 2024; Nain et al. 2024; Kathirithamby et al. 2025; Kim et al. 2025). A suite of unusual genomic features, including exceptionally small genome sizes, sex-specific patterns of endoreduplication, and elevated rates of molecular evolution have been found in Strepsiptera (Johnston et al. 2004; McMahon et al. 2011; Castaño et al. 2024; Kathirithamby et al. 2025). These patterns are thought to reflect selection for replicative and metabolic efficiency in parasitic lineages, along with substantial gene loss accompanying the morphological simplification of obligate endoparasitism (Johnston et al. 2004; Haag et al. 2014; Jackson 2015; Zverkov et al. 2019; Castaño et al. 2024). Collectively, the combination of miniaturized genomes, extreme sexual dimorphism, and diverse host-driven selective pressure makes Strepsiptera a compelling system for studying the evolutionary forces shaping genome content and architecture.

High quality long-read genome assemblies remain rare for Strepsiptera, however, and are currently restricted to two species that stylopize Hymenoptera: *Xenos peckii* (Xenidae), a parasite of social paper wasps (Hymenoptera: Vespidae) (Castaño et al. 2024), and *Stylops aterrimus* (Stylopidae), a parasite of solitary mining bees (Hymenoptera: Andrenidae) (Kathirithamby et al. 2025). These two families account for the vast majority of existing behavioral, physiological, taxonomic, and genomic knowledge of the order, largely because of the abundance and visibility of their hymenopteran hosts (Giusti et al. 2007; Hrabar et al. 2014; Jůzová et al. 2015; Kathirithamby et al. 2015; James et al. 2016; Balzer and Davis 2021; Benda et al. 2021; Millena et al. 2022; Weingardt et al. 2023; Jandausch et al. 2024; Millena et al. 2024). As a result, current genomic understanding of Strepsiptera is disproportionately shaped by lineages parasitizing a single host order, leaving unresolved which genomic signatures reflect broad, order-wide patterns of endoparasitism versus host-specific adaptation to bees and wasps. We hypothesize that co-evolution with specific insect hosts and their distinct life histories may influence the genome evolution of strepsipteran parasites. Alternatively, Strepsiptera may have a universal “skeleton key” strategy enabling successful infection regardless of host, or have evolved specialized post-translational modifications explaining the broad range of insects they parasitize. Regardless, a broader sampling of strepsipteran species is needed to determine if differences in host use are reflected in their genomic makeup.

Within Strepsiptera, the family Elenchidae offers a critical opportunity to address these gaps in knowledge. Rather than infecting social Hymenoptera as seen in the more common Xenidae or Stylopidae, Elenchidae stylopizes solitary planthoppers (Hemiptera: Delphacidae) and remains almost entirely understudied at the genomic level relative to Xenidae and Stylopidae (Millena et al. 2025). Previous work on Elenchidae has been restricted to its morphology, physiology, and taxonomy (Eaton 1892; Fox 1967; Brailovsky 1981; Kathirithamby 1989; Luna de Carvalho 1990; Pohl 1993), developmental biology (Kathirithamby 1983; Kathirithamby et al. 1984; Kathirithamby et al. 1992; Gu et al. 1994; Büning 1998; Carcupino et al. 1998; Maeta et al. 2007; Suraksakul et al. 2022), or ecology, especially in the context of host use (Chiu 1979; Kathirithamby 1979; Stiling and Strong 1982; Hachiya 1988; Stiling et al. 1991; Olmi 1998; Noda et al. 2001). Molecular sampling of Elenchidae has been limited to one or two Sanger sequenced genes for the purpose of barcoding in population ecology or phylogenetic studies (Matsumoto et al. 2011; McMahon et al. 2011). One species of the family, *Elenchus koebelei* Pierce (Figure 1A-B), is a minute (<1.5 mm) crepuscular strepsipteran found parasitizing delphacid planthoppers (Figure 1C) feeding on smooth cordgrass in southeastern U.S. salt marshes (Stiling and Strong 1982; James and Strong 2018). This genus has been the subject of multiple studies thanks to its relative abundance and impact on planthopper populations (Muir 1906; Chiu 1979; Stiling and Strong 1982; James and Strong 2018). Unlike the largely diurnal Xenidae and Stylopidae, adult male *E. koebelei* are active only briefly around dawn (Straka et al. 2011; Hrabar et al. 2014; James and Strong 2018). Recently-developed ultraviolet light trapping methods have enabled reliable capture of adult male *E. koebelei* in the field (James and Strong 2018), providing the specimens used in the present study and ensuring new opportunities to investigate the genomic and physiological specializations for both Elenchidae and Strepsiptera as a whole.

**Figure 1.**
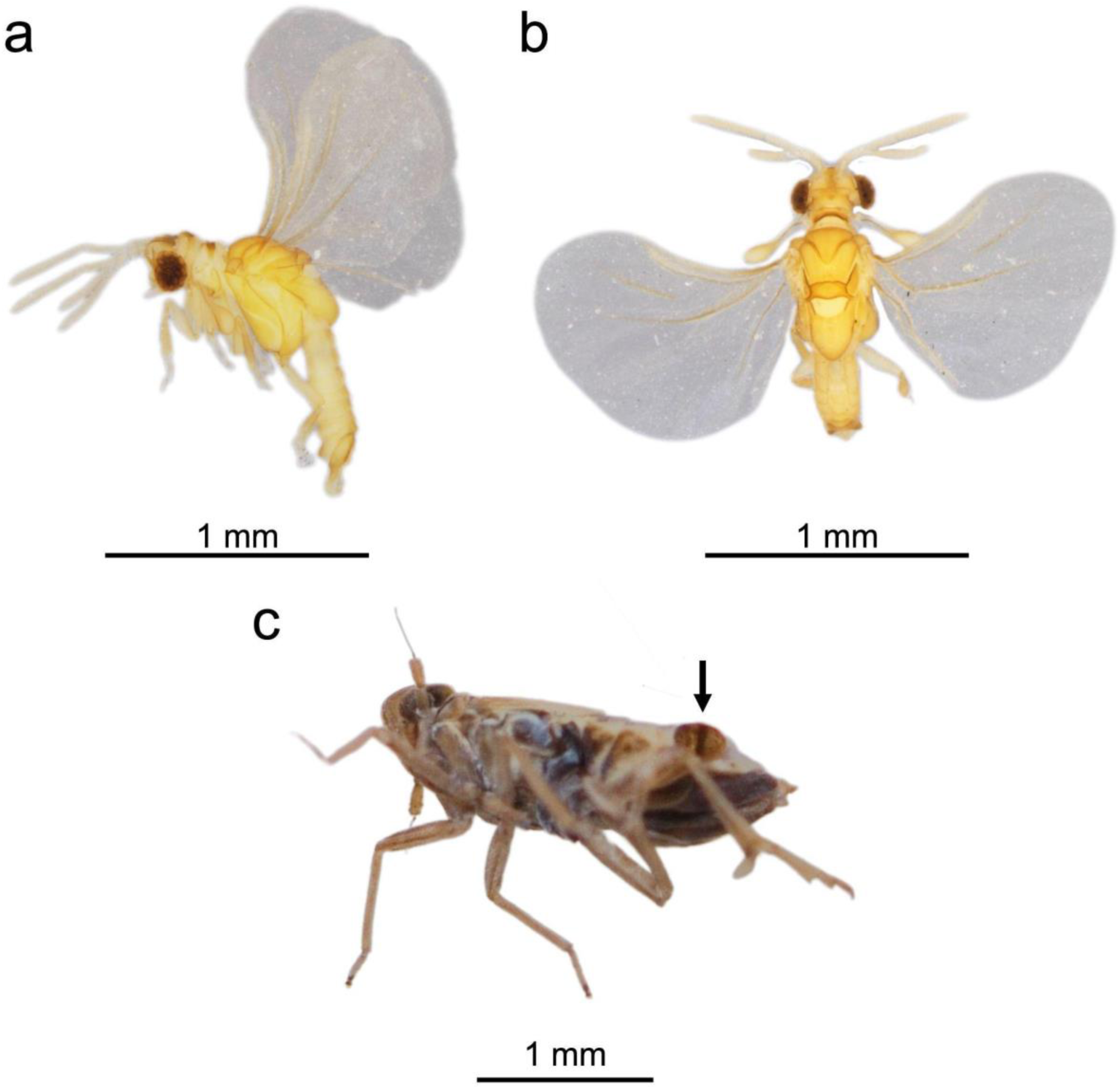
*Elenchus koebelei* male specimen from the Bohart Museum of Entomology (University of California, Davis) collections. **A)** Left side view. **B)** Dorsal view. **C)** Left-ventral view of parasitized *Prokelisia sp.* host nymph, with arrow indicating empty male pupal casing of *E. koebelei*. Image originally from James and Strong (2018).

Here, we present a de novo genome assembly of *Elenchus koebelei* (Figure 1), generated using PacBio Hifi long-read sequencing from adult males collected via ultraviolet light trapping in salt marsh habitat in the southeastern United States. This represents the first NGS genome assembly for a non-Hymenoptera-parasitizing strepsipteran, and is only the third high-quality (as defined by Chain et al. 2009) Strepsiptera genome published to date. The final assembly is comprised of 96 contigs, with a size of 63.7 Mb, N50 of 1.60 Mb, and 32.7% GC content (Table 2). This genome expands the breadth of genomic resources available for Strepsiptera, providing a critical comparative dataset for examining patterns of evolution and biological processes among lineages with divergent host associations. By comparing three genome assemblies representative of different strategies in strepsipteran host use, we investigated the degree to which host biology and infection strategy dictate genomic evolution in this enigmatic order.

**Table 1.**
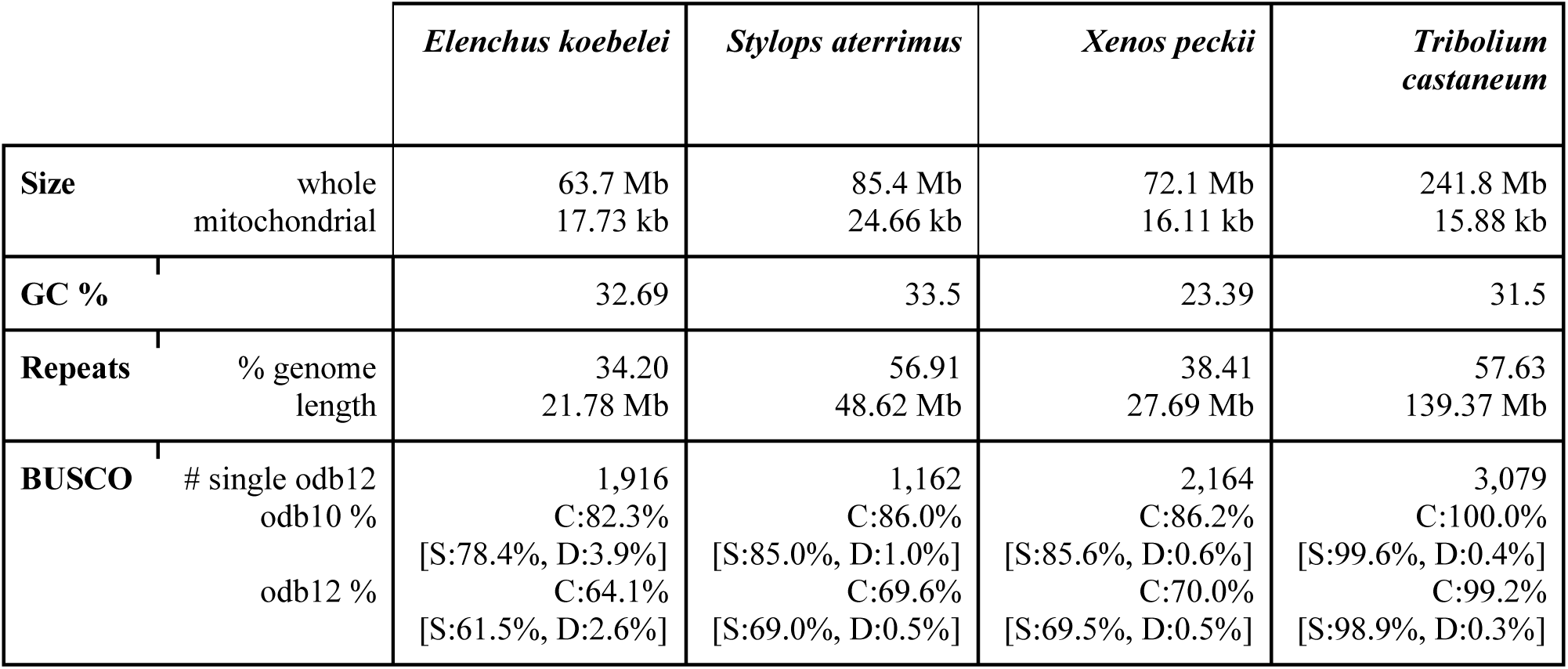
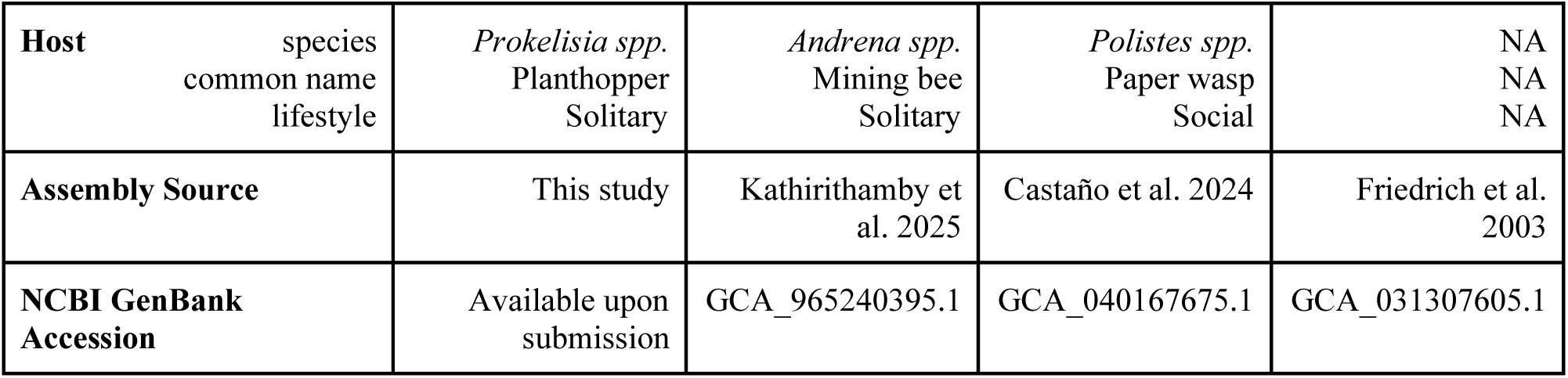
Comparison of biological information and genomic attributes in NGS assemblies between strepsipteran species and a beetle outgroup (*Tribolium castaneum*). For standardization and comparison in this study, repeat content was newly assessed with the Earl Grey pipeline for *Stylops aterrimus* and *Tribolium castaneum*, and BUSCO scores were rerun with odb12 for each species. All other entries are sourced from material published with the respective assemblies. BUSCO abbreviations are as follows: C, complete; S, single-copy; D, duplicated.

**Table 2.** Assembly statistics for NGS strepsipteran genome assemblies. Number of genes for *Stylops aterrimus* was newly determined via annotation with BRAKER3, as the original assembly was not originally annotated. All other entries are sourced from material published with the original assemblies.

| Measure | *Elenchus koebelei* | *Stylops aterrimus* | *Xenos peckii* |
| --- | --- | --- | --- |
| Assembly level | Contig | Chromosome | Contig |
| # contigs | 96 | 179 | 227 |
| N50 | 1.6 Mb | 8.4 Mb | 7.4 Mb |
| L50 | 10 | 3 | 5 |
| Coverage | 86x | 281x | 112x |
| # genes | 8196 | 8974 | 8759 |
| % BUSCO missing (odb10) | 15.8 | 12.3 | 12.1 |
| Sequence tech | PacBio HiFi | PacBio HiFi + Hi-C | PacBio HiFi |

## Methods

### Specimen preparation, DNA extraction, library preparation, and sequencing

We sequenced the genomes of two individuals of *E. koebelei*. Specimens were adult males collected in September 2013 at a salt marsh site in Guana Tolomato Matanzas National Estuarine Research Reserve, Florida, USA (30.0702°N, 81.3447°W). Specific methods for specimen collection and transport are detailed in James and Strong 2018. Following collection, the specimens were stored in molecular grade ethanol in a –80°C freezer at the University of California, Davis Genome and Biomedical Sciences Facility. High molecular weight DNA was extracted from each individual using a Qiagen genomic tip DNA extraction kit (as in Tolman et al. 2023); QIAGEN, Valencia, CA, USA). Because the specimens were minute, we were unable to extract enough high molecular weight DNA for a standard PacBio HiFi library. Instead, we used the PacBio HiFi Ultra-Low library preparation protocol to generate two sequencing libraries from single individuals. The Ultra-Low libraries were barcoded and pooled, then sequenced on a PacBio Revio SMRT cell in CCS mode at the Brigham Young University DNA Sequencing center. We followed the PacBio guidelines for Ultra-Low libraries, including demultiplexing and trimming barcodes with Lima (Biosciences 2023a), and removing PCR duplicate reads with pbmarkdup (Biosciences 2023b).

### Genome assembly and quality assessment

We chose programs and parameters for assembly and optimization in accordance with the methods and findings detailed in Castaño et al. 2024, thus ensuring that our genome assembly would be comparable to the PacBio HiFi genome assemblies for *Xenos peckii* and *Stylops aterrimus*. Prior to assembly, k-mer frequency profiles were generated from the raw HiFi reads using KMC v3.2.1 (Deorowicz et al. 2015) (k=21) to estimate genome size, heterozygosity, and repeat content via GenomeScope 2.0 (Vurture et al. 2017). We then assembled the trimmed, deduplicated reads with Hifiasm v0.20.0 (Cheng et al. 2021) in standard HiFi-only mode, producing primary and haplotype-resolved contig draft assemblies. Assembly contiguity and basic statistics were assessed using QUAST v5.3.0 (Gurevich et al. 2013). Genome completeness was evaluated using BUSCO v6.1.0 (Seppey et al. 2019) against the Insecta OrthoDBv10 (Kriventseva et al. 2019) and OrthoDBv12 (Tegenfeldt et al. 2025) lineage datasets in genome mode, and with Meryl v1.4.1 and Merqury v1.4.1 to assess completeness based on k-mers (Rhie et al. 2020). We proceeded to complete all other analyses with a single primary assembly chosen based on its higher contiguity and gene completeness compared to that of the second *E. koebelei* individual. We included the genome of *Tribolium castaneum* as a reference-quality genome comparison, since it is a model organism belonging to Coleoptera, the sister group of Strepsiptera, and can therefore provide phylogenetically informative context (Millena et al. 2025).

### Genome filtering and annotation

We performed contamination screening and filtering via taxon-annotated GC-coverage plots generated with BlobTools v4.5.3 (Laetsch and Blaxter 2017). To model and mask repeats in the assemblies, we used RepeatModeler and RepeatMasker within the Earl Grey v7.3.0 pipeline (Flynn et al. 2020; Baril et al. 2024). To annotate the genome, we used the BRAKER3 pipeline (Gabriel et al. 2024) with EPmode for GeneMark-EP+ (Brůna et al. 2020), using the Arthropoda database of protein families from OrthoDBv10 as evidence for AUGUSTUS (Stanke and Morgenstern 2005). Additionally, we predicted functional annotations, orthologs, and domains from the BRAKER3 output using eggNOG 5.0 and eggNOG-mapper v2.1.15 (Huerta-Cepas et al. 2019; Cantalapiedra et al. 2021).

### Mitochondrial genome assembly

Using the MitoHifi pipeline (Uliano-Silva et al. 2023), we performed the mitochondrial sequence assembly of *Elenchus koebelei*. For the reference-guided assembly, we chose the mitochondrial genome of *Xenos yangi* (NCBI Accession OK329871) as the most complete and related reference sequence option. We then used MitoHifi to assemble contigs of the mitogenome from our raw HiFi reads (flag -r) and conducted an additional annotation with MITOS2 (Bernt et al. 2013). To verify the content and identity of this assembly, we used McScanX (Wang et al. 2012) to conduct a syntenic analysis with the other available mitogenomes of Strepsiptera: *Dipterophagus daci* (NCBI accession MW233588.1), *Mengenilla moldrzyki* (JQ398619.1), *Stylops aterrimus* (OZ251264.1), *Xenos peckii* (CM079275.1), and *Xenos yangi* (OK329871.1). We anchored the beginning of each mitochondrial sequence at the cytochrome c oxidase subunit I (COI) gene for gene order comparison, and visualized the results with the *circlize* package in R (Gu 2013).

### BUSCO gene comparisons

To perform missing gene comparisons, we ran BUSCO on all three parasite species against both Insecta OrthoDBv12 (odb12) and v10 (odb10). We specifically performed these analyses because gene number and percentage completeness vary significantly between the two versions, and both the *Xenos peckii* and *Stylops aterrimus* assemblies were assessed for completeness with odb10 in their original publications (Castaño et al. 2024; Kathirithamby et al. 2025). We also included *Tribolium castaneum* in this analysis to determine which losses were likely to be specific to the order. To confirm BUSCOs were truly missing from the assemblies, we used BLAST (Camacho et al. 2009) to align each missing BUSCO sequence against the raw reads. To evaluate which genes were truly missing from *Elenchus koebelei*, and not just an artefact of misassembly, we considered only BUSCOs that were listed as absent in all *E. koebelei* assemblies (primary, alternate haplotype, and the assemblies from the second sample) as missing from the species. We then compared the missing gene lists using the command line tools sort, grep, awk, and comm, and evaluated the function of genes of interest by querying OrthoDB orthogroup descriptions for the BUSCO IDs. We conducted and visualized functional enrichment tests on the eggNOG annotations for each BUSCO gene set using the R packages *funfea* (Charest et al. 2025), *clusterprofiler* (Wu et al. 2021), *GO.db* (Carlson et al. 2019), and *KEGGREST* (Tenenbaum et al. 2026). For these enrichment analyses, we treated all BUSCO IDs in the insecta_odb12 lineage as our background gene set.

## Results and Discussion

### Sequencing

PacBio HiFi sequencing of the primary *Elenchus koebelei* specimen generated 5.56 Gb of data (Q > 45), comprising 826,916 HiFi reads with mean read length 6,726 bp and an estimated 86x coverage of the genome. The secondary individual yielded a lower-coverage (77x) dataset that was used only for BUSCO cross-validation and comparative missing-gene analysis (see BUSCO comparisons, below), as its assembly was less contiguous than that of the primary specimen. The primary assembly comprised 96 contigs after filtering, and had an N50 of 1.60 Mb, L50 of 10, and BUSCO odb10 score of 82.3%, while the alternative consisted of 2,616 contigs with an N50 of 1.10 Mb, L50 of 20, and BUSCO odb10 score of 64.9%. k-mer profiling (k=21) of the raw HiFi reads with GenomeScope 2.0 for the primary specimen estimated a haploid genome length of 50.52-50.61 Mb with 1.35% heterozygosity and 16.68% duplication (Figure 2). The model converged well (fit 82-94%) with a low estimated read error rate and low heterozygosity, indicating that the k-mer spectrum was well-resolved.

**Figure 2.**
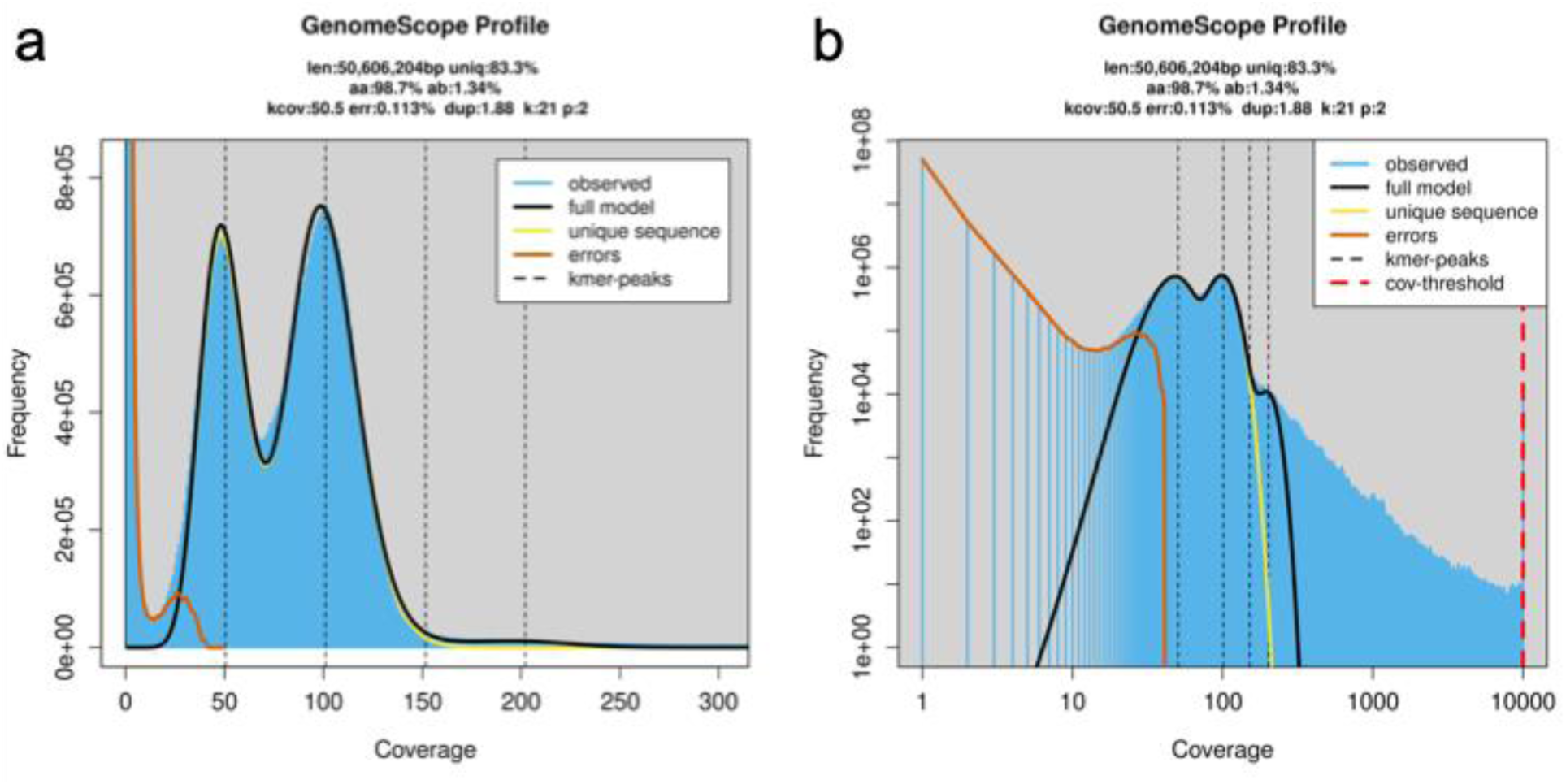
Genomescope2 k-mer (21) distribution from the adapter-trimmed PacBio HiFi raw reads. Plot displays estimation of genome length (len), percentage of the genome that is non-repetitive (uniq), homozygous rate (aa), heterozygous rate (ab), mean k-mer coverage for heterozygous bases (kcov), read error rate (err), average rate of duplications (dup), k-mer size used on run (k), and ploidy (p). **A)** Expected and obtained linear plot distribution of coverage according to the model. **B)** Expected and obtained logarithmic plot distribution of coverage according to the model. The four peaks correspond to the mean coverage levels of the unique heterozygous, unique homozygous, repetitive heterozygous and repetitive homozygous sequences, respectively.

### Assembly statistics

Prior to filtering, the primary *Elenchus koebelei* assembly comprised 147 contigs spanning 64.26 Mb, of which contigs taxonomically annotated as Arthropoda accounted for 62.91 Mb across only 125 contigs (Figure S1). This reflected a highly-fragmented, low coverage tail that is possible for HiFi assemblies of minute specimens prepared with an Ultra-Low library protocol (Schardt and Bálint 2023; Schell et al. 2025). Blobtools filtering revealed 47 < 0.02 Mb contigs that had GC content of less than 20% and 22 non-Arthropoda contigs. We removed the 47 short contigs and 4 non-target contigs belonging to Streptophyta, as none of these contigs influenced BUSCO completeness, and retained contigs annotated as no-hit (read mapping percentage 0.27%, 14 contigs), Mollusca (0.25%, 3 contigs), and Chordata (0.19%, 1 contig) due to contamination status ambiguity. This resulted in a final genome assembly of 96 contigs spanning 63.65 Mb (Figures 3, 4). QUAST output on the filtered assembly reported an N50 of 1.60 Mb (L50 = 10), largest contig of 5.40 Mb, GC content of 32.69%, and zero ambiguous bases, indicating a highly contiguous, gap-free assembly (Table 2).

**Figure 3.**
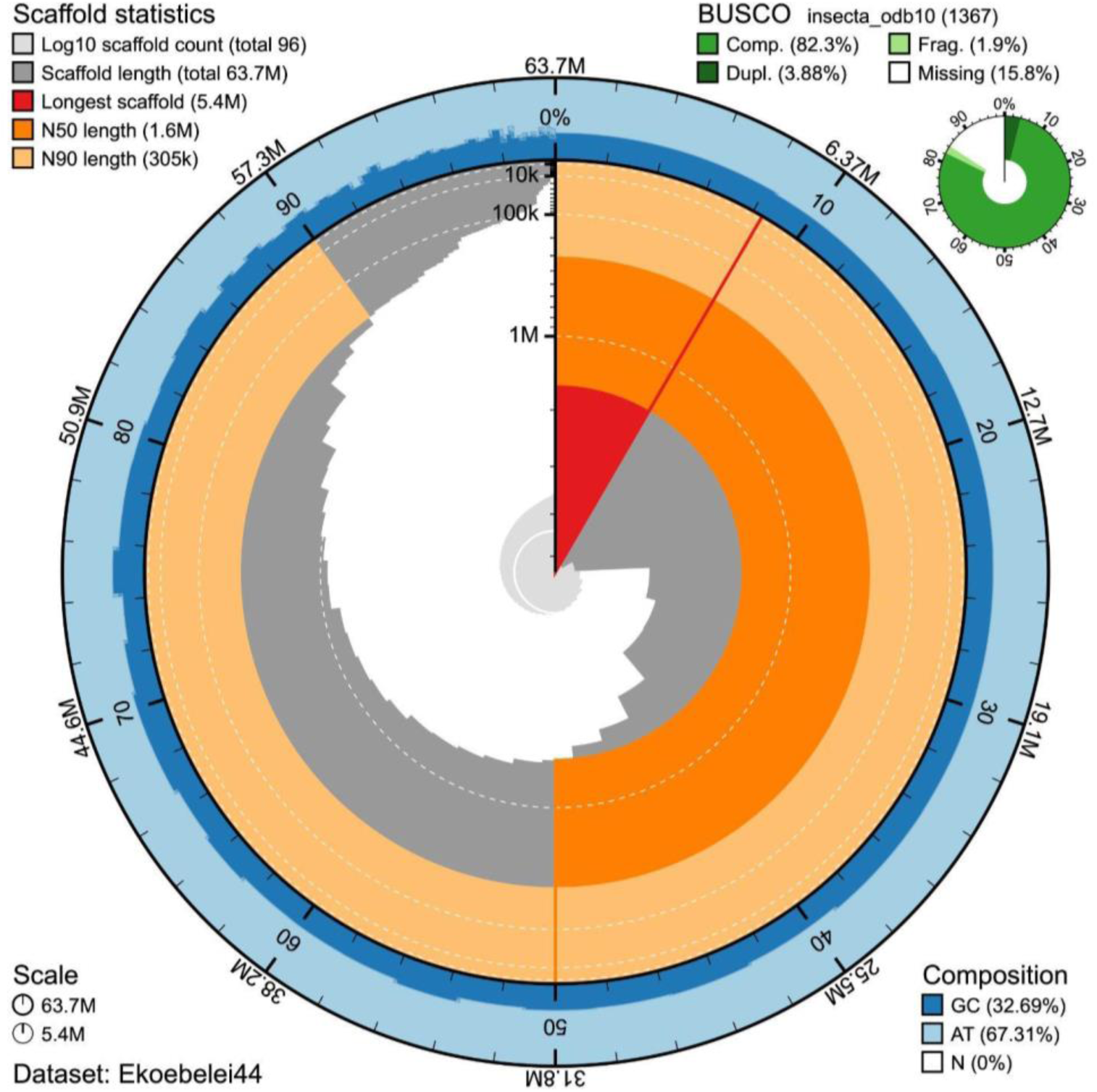
BlobToolKit snail plot visualization of *Elenchus koebelei* genome assembly. Completeness and contiguity of the filtered dataset are plotted as a circle representing the full assembly (63.7 Mb) in 96 contigs. The uniform GC content (32.69%) is plotted in blue as the outside track of the circle. The inner track represents the longest contig (5.4 Mb, highlighted in red), the N50 (1.5 Mb, highlighted in dark orange), and the N90 (305 kb, highlighted in light orange). Lengths of each contig are visible in the height of the gray steps against the vertical radius axis. BUSCO scores (odb10) are in the top right corner in green.

**Figure 4.**
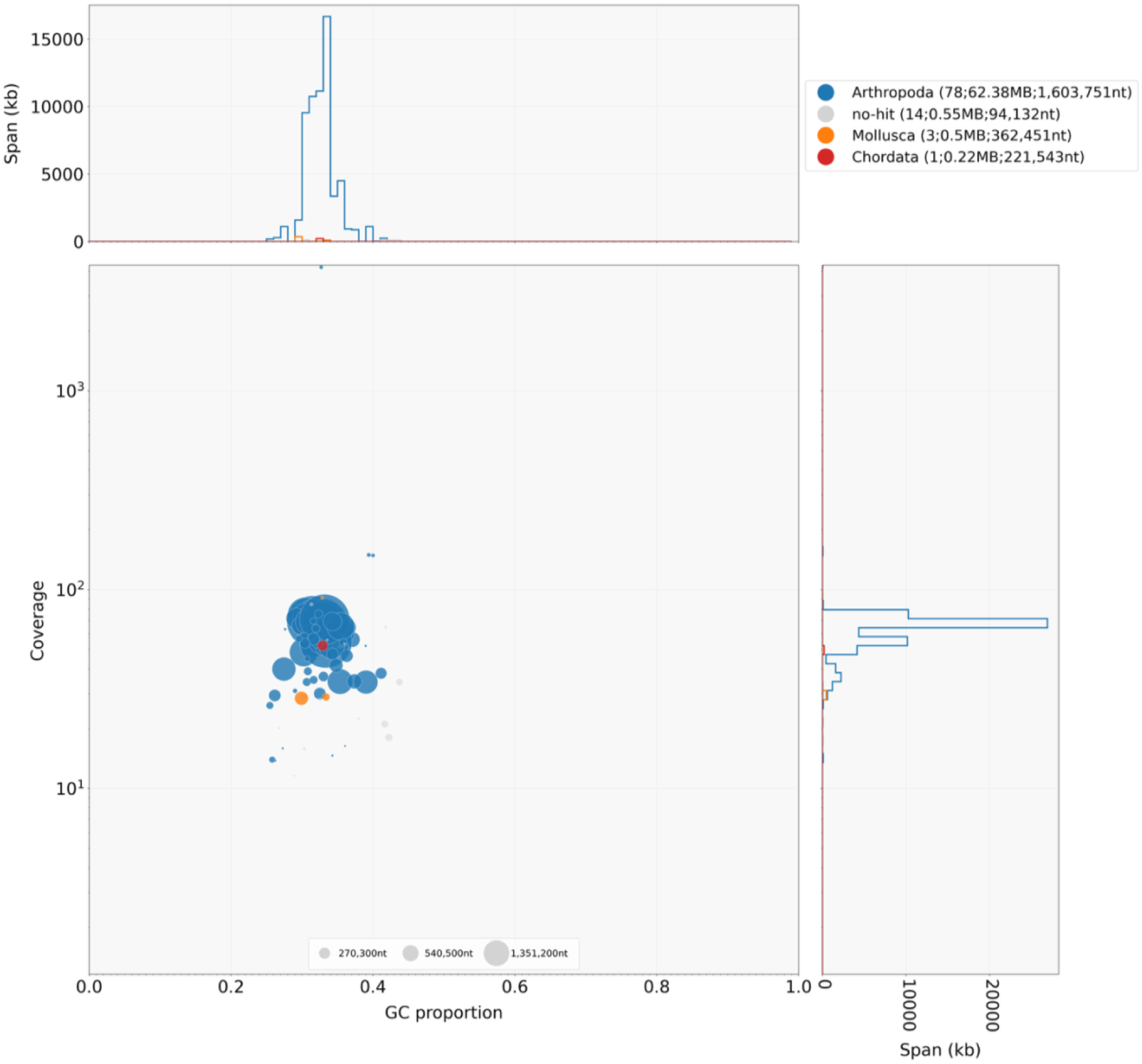
Blobplot representing filtered *Elenchus koebelei* contigs. Contigs are represented as circles, with diameters proportional to sequence length and colored by taxonomic affiliation (blue, Arthropoda; gray, no-hit; orange, Mollusca; red, Chordata). Coverage (right) and GC (top) histograms are weighted by total span of sequences occupying each bin.

### Repeat content

Using Earl Grey, we identified 34.20% (21.78 Mb) of the *E. koebelei* genome as repetitive, dominated by unclassified elements (20.1% of genome), simple/microsatellite repeats (4.6%), DNA transposons (4.1%), LTR retrotransposons (3.9%), and LINEs (1.5%) with SINEs and Rolling Circle elements each contributing < 1% (Figure 5A). The single largest individual repeat family is an unclassified element (RND-1_FAMILY-329#Unknown), contributing 1.61 Mb of genomic coverage across 402 copies and a moderate mean Kimura divergence of 0.18 (range 0-0.55) (Figure 5B). A skew towards unclassified repeat content is also seen in the *Stylops aterrimus* (36.06%, Figure S2) and *Xenos peckii* (27.80%) assemblies. This highlights how poorly the repeat landscape of Strepsiptera is represented in previously-sequenced, existing reference libraries, and presents a substantial need for future curation.

**Figure 5.**
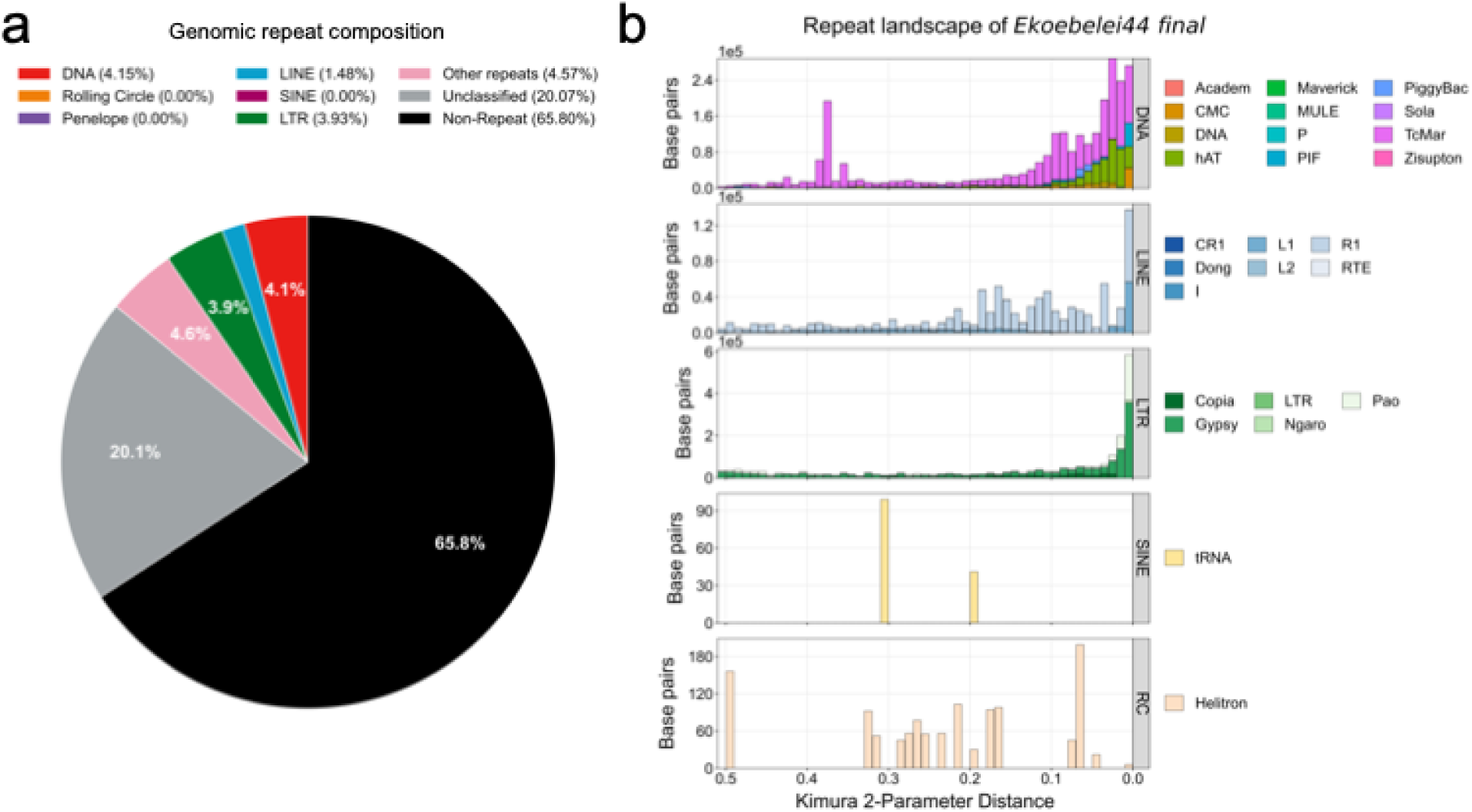
Repeat content classification of the *Elenchus koebelei* genome assembly using the Earl Grey pipeline. **A)** Genomic composition of the genome assembly in TE classifications. **B)** Repeat landscape of the assembly, by repetitive element superfamilies.

### Genomic structure of Strepsiptera

Across the three strepsipteran assemblies, genome size varies more than 30% (63.6-85.4 Mb, Table 1). This may be in part due to repeats, as the non-repetitive content of each assembly falls within a narrower length range (Figure 6). These repetitive elements require further investigation; they may contain informative biological differences, especially with the increase in repetitive content seen in *S. aterrimus*. However, difficulty in assembling repeats can also lead to assembly error in a way that may artificially inflate genome size, especially when sequencing samples from pooled individuals (Futschik and Schlötterer 2010; Tørresen et al. 2019). Regardless, unclassified repetitive elements dominate in the repeat content of all three genomes. *E. koebelei* (solitary planthopper host) and *X. peckii* (social wasp host) are more similar in size and repeat content than *S. aterrimus* (solitary bee host), suggesting genome size variation is not related to the sociality of the host insect. GC content is likewise similar between *E. koebelei* and *S. aterrimus* (32.7% and 33.5%) but notably lower in *X. peckii* (23.4%), and total gene counts fall within a comparatively narrow range (∼8,100-9,000). The broad genomic structure of Strepsiptera therefore seems to be consistent and without a corresponding trend tied to host use. Reduced gene content and variable but generally elevated repeat turnover relative to a free-living beetle representative (Table 1) is consistently shown in all three species. This suggests that host-specific adaptations, if present, are not necessarily encoded in gross genome structure; however, more sampling is needed to confirm this. Specialization may instead be present at the level of specific gene family expansion and contraction, or in post-translational regulation.

**Figure 6.**
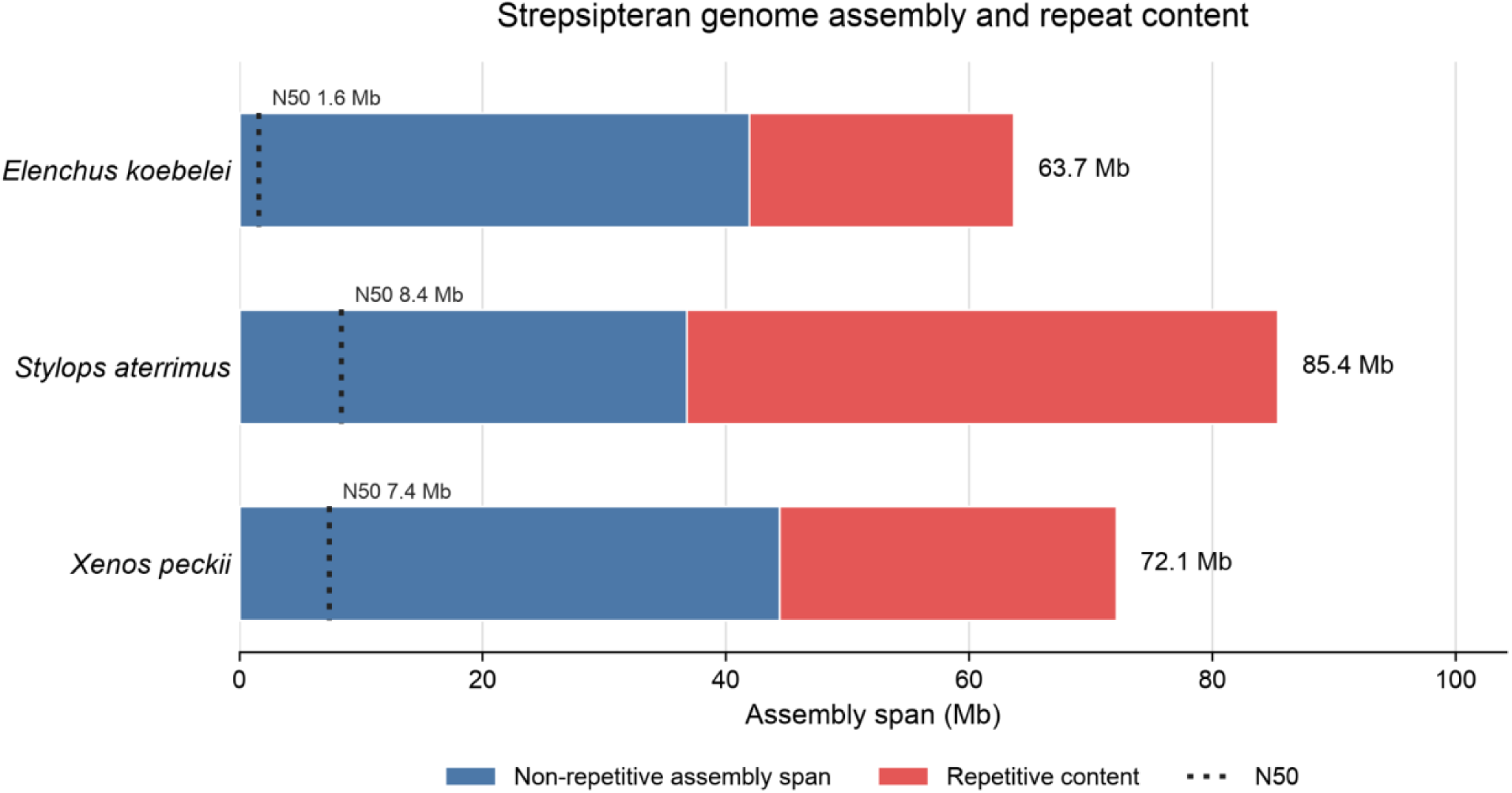
Bar plot representation of select Strepsiptera assembly statistics. Haploid assembly size is in blue, and repetitive content is in red. N50 statistics are visible as dotted lines.

### Mitochondrial genome

The *Elenchus koebelei* mitogenome assembled from our raw reads is 17.73 kb, containing 34 genes representative of the typical metazoan mitochondria (Figure 7A). We recovered and annotated 11 protein-coding genes, a full complement of 22 tRNAs, and a single rRNA. There were three genes not recovered: ATP6, ATP8, and rrnS (16S rRNA). This mitogenome is broadly consistent in gene content and order with previously published Strepsiptera mitogenomes (Figure 7B). McScanX synteny analysis of *E. koebelei* against *Dipterophagus daci*, *Mengenilla moldrzyki*, *Stylops aterrimus*, *Xenos peckii*, and *X. yangi* (Figure 7B) recovered a single, fully collinear block of protein-coding genes between *E. koebelei* and each other strepsipteran mitogenome compared, with 91.46% of all shared genes falling into collinear blocks mitogenome-wide (75 of 82 total genes). Gene order and relative orientation were consistent with *X. peckii* and *X. yangi* (n = 10−12 genes, evalue ≤ 1.1E-11) and with *D. daci* and *M. moldrzyki* (n = 8−11 genes, evalue ≤ 1.4E-08), but weaker with *S. aterrimus* (n = 5, evalue = 0.11) as the only non-significant comparison. Because ATP6 and ATP8 were not recovered in the *E. koebelei* annotation, gene order comparisons here relative to our focal species are limited to the 11 shared protein coding genes. The absence of detectable rearrangement amongst the six strepsipteran mitogenomes analyzed here supports a broadly conserved mitochondrial gene order throughout Strepsiptera, especially given the taxonomic distance between the additional species included in this analysis; *Mengenilla moldrzyki* belongs to the earliest-derived extant family Mengenillidae, which parasitizes silverfish and has free-living females, and *Dipterophagus daci* is in the family Halictophagidae and parasitizes true fruit flies (McMahon et al. 2011; Towett-Kirui et al. 2021). Gene content, however, may be a different story—this assembly is now the second example of MITOS2 being unable to resolve annotation of the 16S rRNA gene in a strepsipteran (Figure 7A), providing additional evidence that this gene has highly divergent regions in these parasites (Towett-Kirui et al. 2022; Castaño et al. 2024).

**Figure 7.**
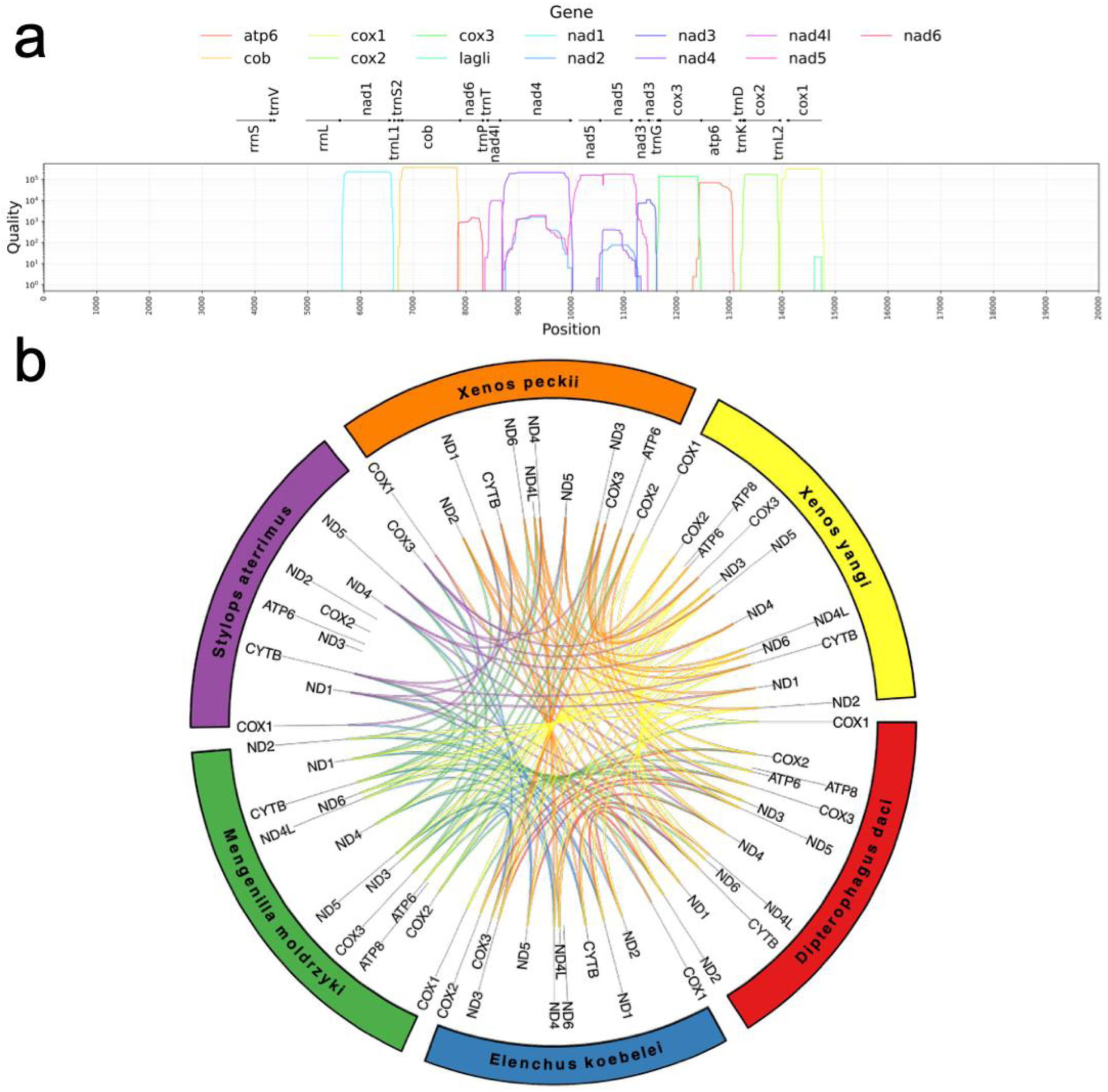
*Elenchus koebelei* mitochondrial genome assembly analyses. **A)** MITOS2 annotation and protein prediction plot. **B)** Strepsiptera mitogenome synteny visualization. All mitogenomes in this synteny plot have been anchored at COX1 for syntenic block detection and order comparison.

### Substantial loss of BUSCO genes within Strepsiptera

In comparison to a reference assembly of the *Tribolium castaneum* genome (missing 19 BUSCOs), 619 BUSCOs were uniquely missing amongst the strepsipterans, and 157 were uniquely absent in the *Elenchus* assembly (Figure 8E). The 619 BUSCOs missing in all three strepsipteran assemblies represent a substantial portion of the genetic toolkits underlying sensory structures, cellular architecture, and developmental complexity. This is consistent with the extreme morphological and ecological specialization exhibited in these obligate endoparasites. A commonality between the sets of missing BUSCOs from all three assemblies was enrichment of GO terms involving intraciliary transport and ciliary structure (Figure 8A–C), consistent with reduced sensory or ciliary investment across the obligately endoparasitic Strepsiptera. Similar loss at a genomic level has been observed in parasitic nematomorphs, with links to a loss of cilia machinery and tentative reduction of sensory genes (Cunha et al. 2023). In this system, however, these terms were only significantly enriched in the individual missing gene sets for *Elenchus* and *Stylops* in this study (e.g., “intraciliary transport,” GO:0042073; in *Elenchus* 18/781 genes, adjusted p = 3.8E–04, in *Stylops* 19/705 genes, adjusted p = 4.25E–06; Figure 8A–B). Though this may imply a possible route of investigation into the molecular differences between strepsipterans infecting solitary vs social hosts, further examination is necessary to definitively test these connections.

**Figure 8.**
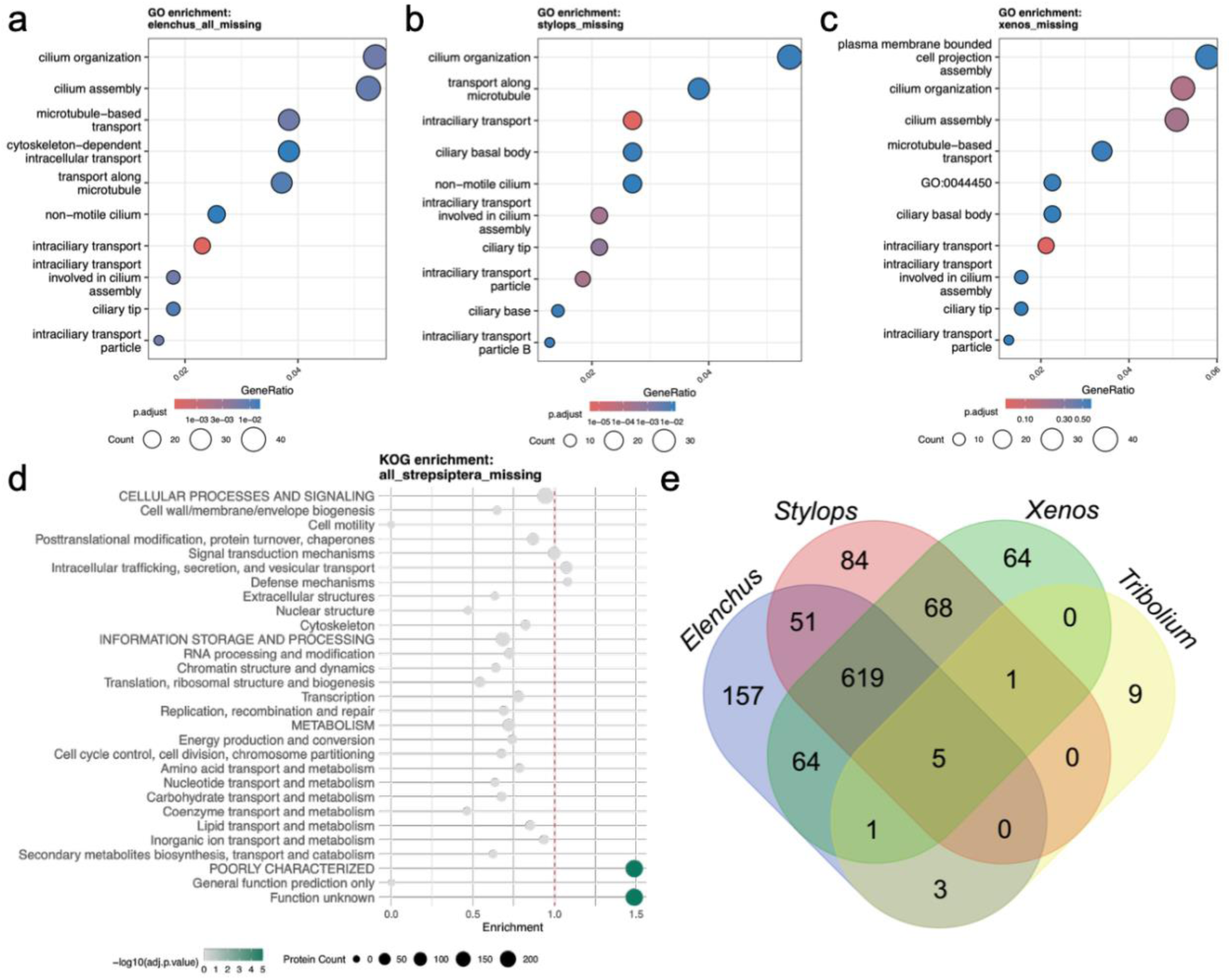
BUSCO enrichment and comparison visualizations. **A)** Lollipop plot for GO term enrichment of missing genes in all *Elenchus koebelei* assemblies included in this study. **B)** Lollipop plot for GO term enrichment of missing genes in the *Stylops aterrimus* genome assembly. **C)** Lollipop plot for GO term enrichment of missing genes in the *Xenos peckii* genome assembly. No terms are significantly enriched (see p.adjust legend scale). **D)** Lollipop plot for KOG term enrichment for the set of missing genes common to all three strepsipteran genome assemblies. **E)** Overlap in BUSCO IDs (odb12) missing from each of the three strepsipteran species assemblies and one beetle outgroup (*Tribolium*).

KEGG pathway enrichment for genes missing specifically in the *E. koebelei* assembly was comparatively weak, with only nominal enrichment (p > 0.01, no terms passing FDR significance correction) for neuromodulary and developmental pathways such as cholinergic and dopaminergic synapse signaling. This pattern was interesting given a potential tie to the compressed adult activity window in Elenchidae (crepuscular vs. diurnal in Xenidae and Stylopidae), though this remains speculative without functional validation. Notably, KOG enrichment analyses returned “POORLY CHARACTERIZED: Function unknown” as the largest category in the overall Strepsiptera missing-BUSCO gene set (234/1815 proteins, adjusted p = 2.91E–09; Figure 8D), emphasizing how much of strepsipteran gene loss is not yet mechanistically interpretable.

## Conclusion

Our results indicate that Strepsiptera possesses a broadly conserved genomic blueprint for obligate endoparasitism that is largely independent of host order or host social structure. Genome size, GC content, gene number, and mitochondrial gene order all vary within a narrow, comparable range across the *Elenchus koebelei*, *Stylops aterrimus*, and *Xenos peckii* assemblies. The genes lost across all three species converge on shared functional categories rather than host-specific signatures. This favors a universal model of strepsipteran genomic specialization for parasitism over one requiring host-specific genomic tuning, though we leave open the possibility that such tuning occurs within gene family copy number, expression, or post-translational regulation. These prospective explanations are compelling prompts for future studies. Future work with additional sampling of genome assemblies and transcriptomes across a broader sampling of strepsipteran species and corresponding host associations are vital for answering these questions in greater depth. As the first genome assembly for a parasite of its kind, the *E. koebelei* genome and the analyses presented here provide a foundation for future comparative work.

## Data Availability

The primary *de novo* whole genome PacBio HiFi assembly was deposited in NCBI under the BioProject accession PRJNA1427140. Pseudohaplotypes for primary and secondary samples are available under BioProject accessions PRJNA1529240–3. The raw genomic sequencing reads were deposited under the NCBI Sequence Read Archive under the accession number SAMN55904791. Sample-specific SRA numbers are SRR37329079 (primary) and SRR37329080. The GFF file with ab initio gene predictions from BRAKER3 and the BUSCO gene sets are available in Dryad.

## Acknowledgments

This work is dedicated to the late Edward Wilcox in honor of his work at the BYU DNA Sequencing Facility. The University of California, Davis and David Coil provided storage and access to the frozen specimens collected by M.J. and sequenced in this study. Special thanks to Maria Castaño, Ethan Tolman, Juan Martín Ferro, the BYU SPIN Workshop, Amanda Larracuente, and members of the Trop Bio Lab at the University of Rochester for scripts and guidance on editing or analyses. Thanks also to members of the Ware Lab at the AMNH for logistical support.

## Author Contributions

R.J.M. – formal analysis, investigation (primary), visualization, data curation, writing - original draft (primary)

P.B.F. – methodology, resources, funding acquisition, software, validation, writing - review and editing

M.J. – resources, writing - review and editing

E.R.T. – validation, data curation, writing - review and editing

F.M.K.U. – conceptualization, resources, methodology, writing - review and editing, supervision

J.L.W. – conceptualization, project administration, funding acquisition, resources, writing - review and editing, supervision

---

**Figure S1.**
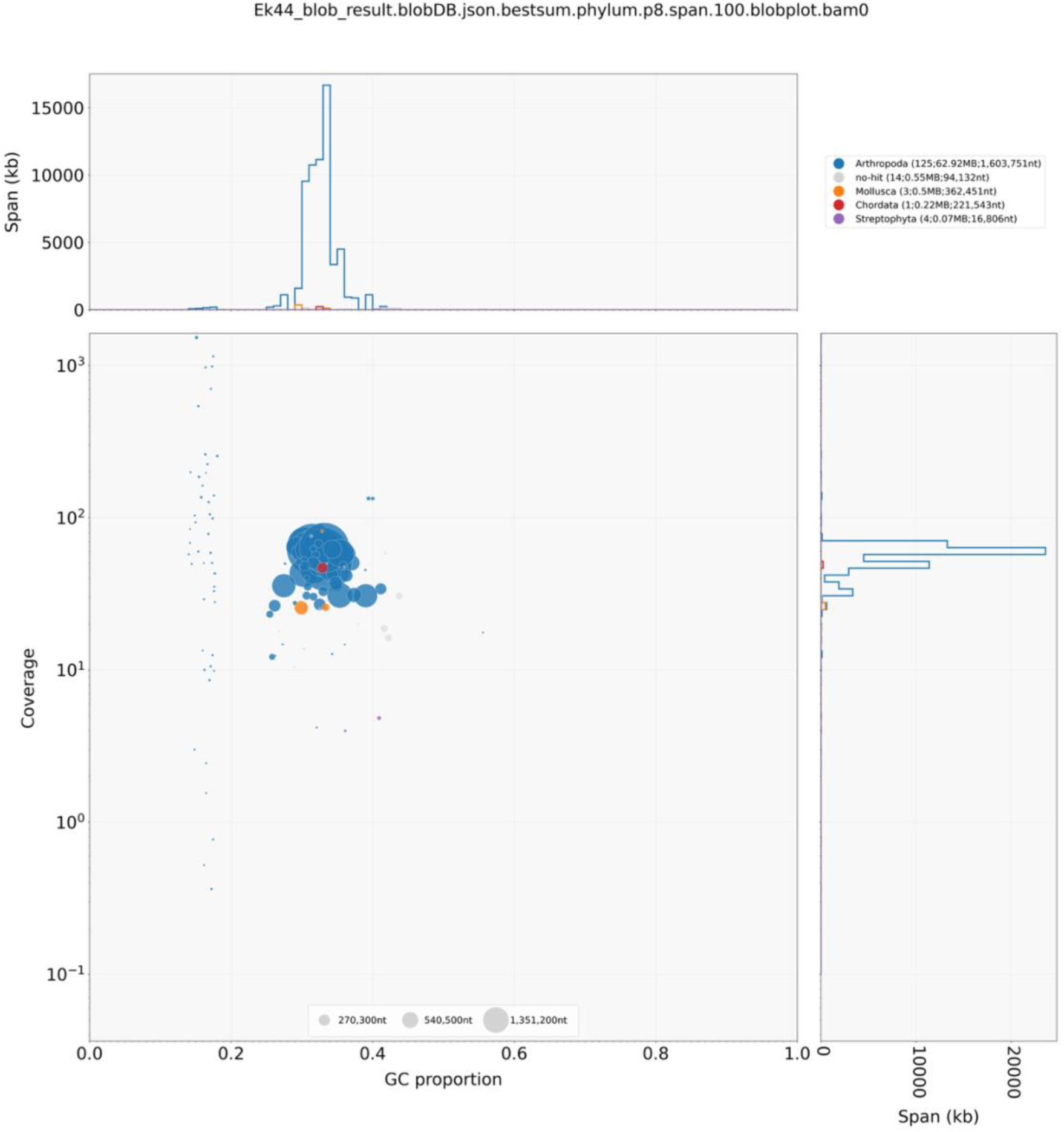
Blobplot representing unfiltered *Elenchus koebelei* contigs. Contigs are represented as circles, with diameters proportional to sequence length and colored by taxonomic affiliation (blue, Arthropoda; gray, no-hit; orange, Mollusca; red, Chordata). Coverage (right) and GC (top) histograms are weighted by total span of sequences occupying each bin.

**Figure S2.**
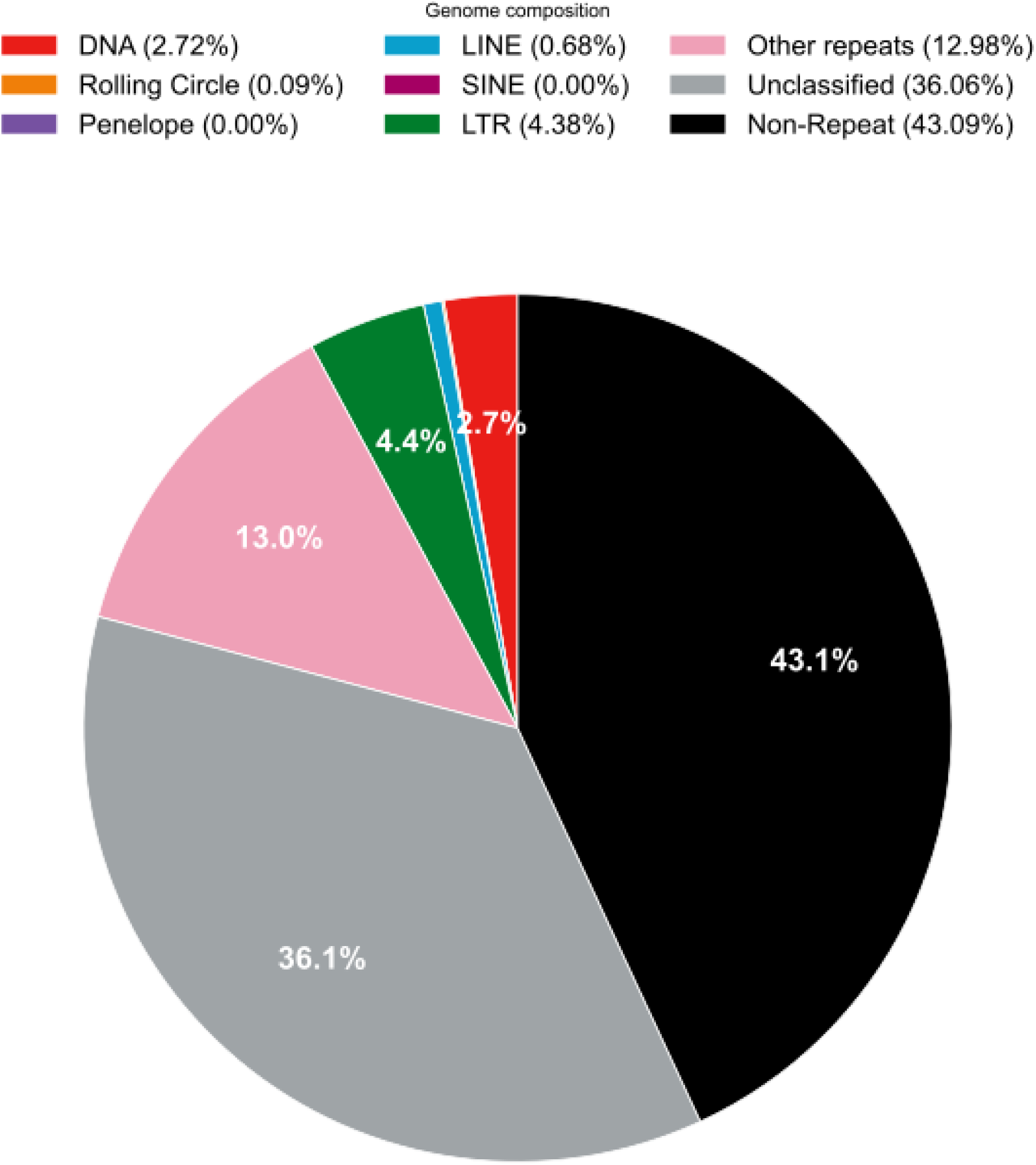
Repetitive element content of *Stylops aterrimus*, generated by the Earl Grey pipeline.

